# Phytochemical Content and Antimicrobial Activity of *Aloe Vera* extract of Marathwada Region, India

**DOI:** 10.1101/2025.06.17.660060

**Authors:** M. Nagargoje, N. Chaudhari, S. N Harke, A. S Khemnar, S. K Barik

## Abstract

Aloe vera (*Aloe barbadensis*) is widely recognized for its therapeutic properties due to its rich phytochemical composition, including proteins, carbohydrates, and steroids. The study aimed to evaluate the efficiency of solvents such as ethanol and methanol for extracting bioactive compounds and to analyse the phytochemical contents of Aloe vera extracts using various biochemical assays. The dinitro salicylic acid test (DNSA) quantified reducing sugars with ethanol extracts yielding 870 mg/ml compared to 450 mg/ml for methanol extracts. The Biuret test for proteins revealed concentrations of 13.8 mg/ml in ethanol extracts and 13.03 mg/ml in methanol extracts. The Barfoed’s test indicated carbohydrate concentrations of 960 mg/ml in ethanol extracts. A qualitative test for steroids confirmed their presence in both solvents, with ethanol showing a more intense reaction, suggesting a higher concentration of steroidal compounds. The results establish ethanol as a more efficient solvent for extracting phytochemicals from Aloe vera. Antimicrobial activity test against E. coli demonstrated superior inhibition zones for ethanol extract, highlighting their potential as natural antibacterial agents. The research indicates the significance of solvent selection in optimizing the therapeutic utility of Aloe vera. It provides a foundation for further exploration into its bioactive compounds and mode of action.

## 1. Introduction

Aloe vera, scientifically known as *Aloe barbadensis*, is a succulent plant belonging to the *Asphodelaceae* family. Renowned for its thick, fleshy leaves filled with a clear gel, Aloe vera contains phytochemicals, including polysaccharides, glycoproteins, amino acids, vitamins (such as A, C, and E), and calcium and magnesium. These phytochemical compounds contribute to antioxidant, anti-inflammatory, and immunomodulatory properties. Consequently, Aloe vera has been widely utilized in both traditional medicine and modern healthcare for treating various ailments [1].

Several biochemical tests can be employed to assess its phytochemical composition of Aloe vera. The DNSA test quantifies reducing sugars present in Aloe vera extracts by measuring optical density at 670 nm after reacting the sample with dinitro salicylic acid (DNSA). This allows researchers to determine the amount of reducing sugars that may contribute to the overall health benefits of Aloe vera. The Biuret test detects proteins and peptides in the extracts by adding a biuret reagent that reacts with peptide bonds to produce a colour change. The intensity of this colour measured at 580 nm correlates with protein concentration which is essential for various biological functions. Additionally, Barfoed’s test differentiates between monosaccharides and disaccharides through a colour change observed after heating with Barfoed’s reagent [2,3]. A qualitative test could identified steroid compounds by mixing the extract with concentrated sulphuric acid and chloroform. The formation of a distinct ring indicates their presence and contributes to Aloe Vera’s anti-inflammatory and immunomodulatory effects [4].

The Aloe vera extraction is significantly influenced by the choice of solvent used during the extraction process. The study compares two common solvent systems: ethanol and methanol. Ethanol is often preferred for its safety profile and effectiveness in extracting a broad spectrum of polar and non-polar compounds while being suitable for human consumption. In contrast, methanol is more polar and may extract different profiles of phytochemicals but raises concerns regarding toxicity and is generally not recommended for food or cosmetic applications. The research aims to identify the solvent system in a better concentration of beneficial compounds from Aloe vera. Understanding the phytochemical composition of Aloe vera extracts is essential for maximizing their therapeutic potential. The research seeks to determine which extraction method optimizes the yield of bioactive compounds that contribute to Aloe vera’s antimicrobial properties.

The findings could inform future applications in pharmaceuticals and natural remedies, enhancing the effectiveness of Aloe vera in treating infections and promoting overall health while addressing challenges such as antibiotic resistance in contemporary medicine [5].

The persistent global challenge of antimicrobial resistance has catalysed scientific exploration of botanical alternatives with Aloe vera emerging as a promising Phyto therapeutic intervention. The systematic investigation comprehensively evaluates the antimicrobial efficacy of Aloe vera extract against Escherichia coli, employing sophisticated extraction methodologies to elucidate bioactive compound profiles [6]. Thus, we focused our research on phytochemical analysis and antimicrobial analysis of region-specific Aloe vera (*Aloe barbadensis*) of Chhatrapati Sambhajinagar, Maharashtra, India.

## 2. Material and Method

The Aloe vera plant is used as a source of material for the study. The various biochemical tests are performed to know about the concentration of proteins, carbohydrates and steroids.

### 2.1 Preparation of Plant Extract

The air-dried Aloe vera plant materials were powdered using a mechanical grinder. The dried plant powder sample was extracted with 99.5% methanol/Ethanol three times by the cold maceration method. The sample was kept in rotatory shaker for 2 days. The crude extracts were filtered with filter paper [4]. The Aloe vera extract prepared is given in Fig-1 (a) *Aloe Vera* was washed thoroughly (b) Chopped it into pieces (c) Air dried *aloe* was powdered (d) Powder was extracted

**Fig 1.**
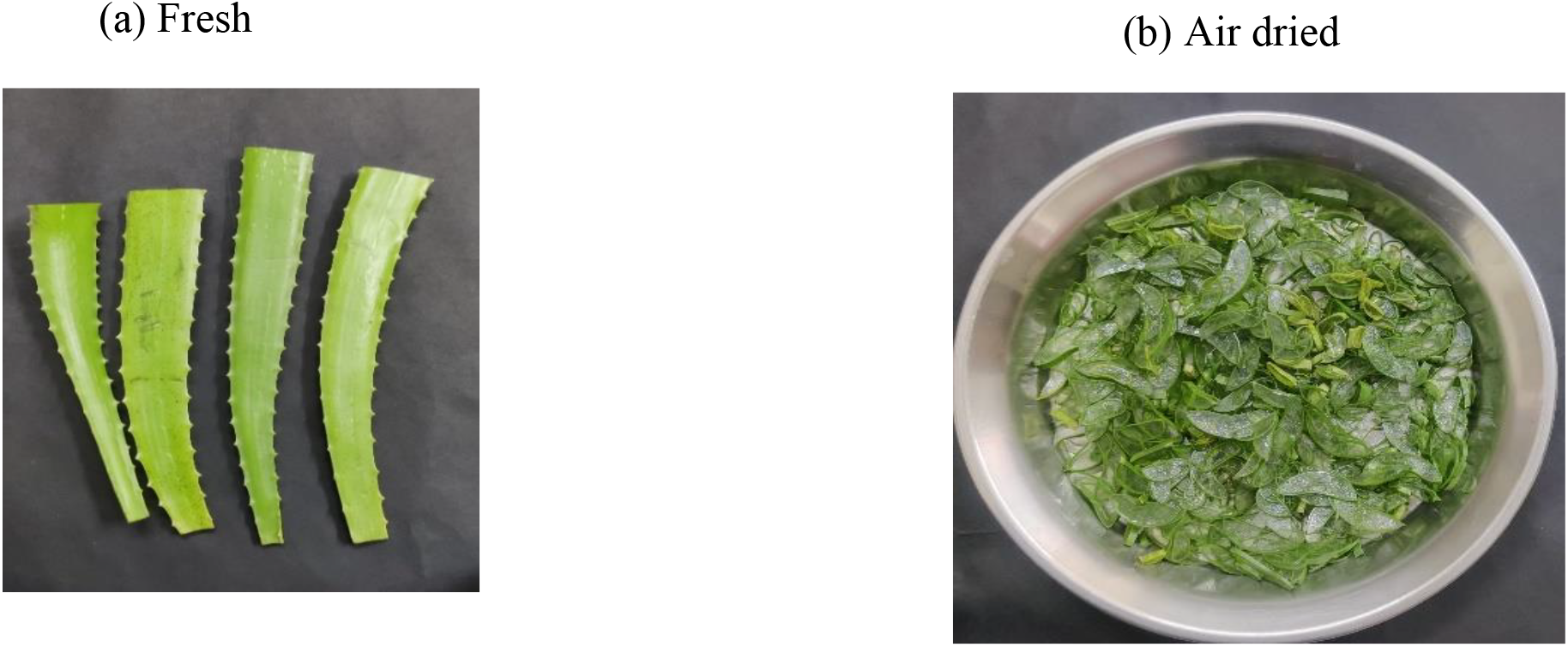
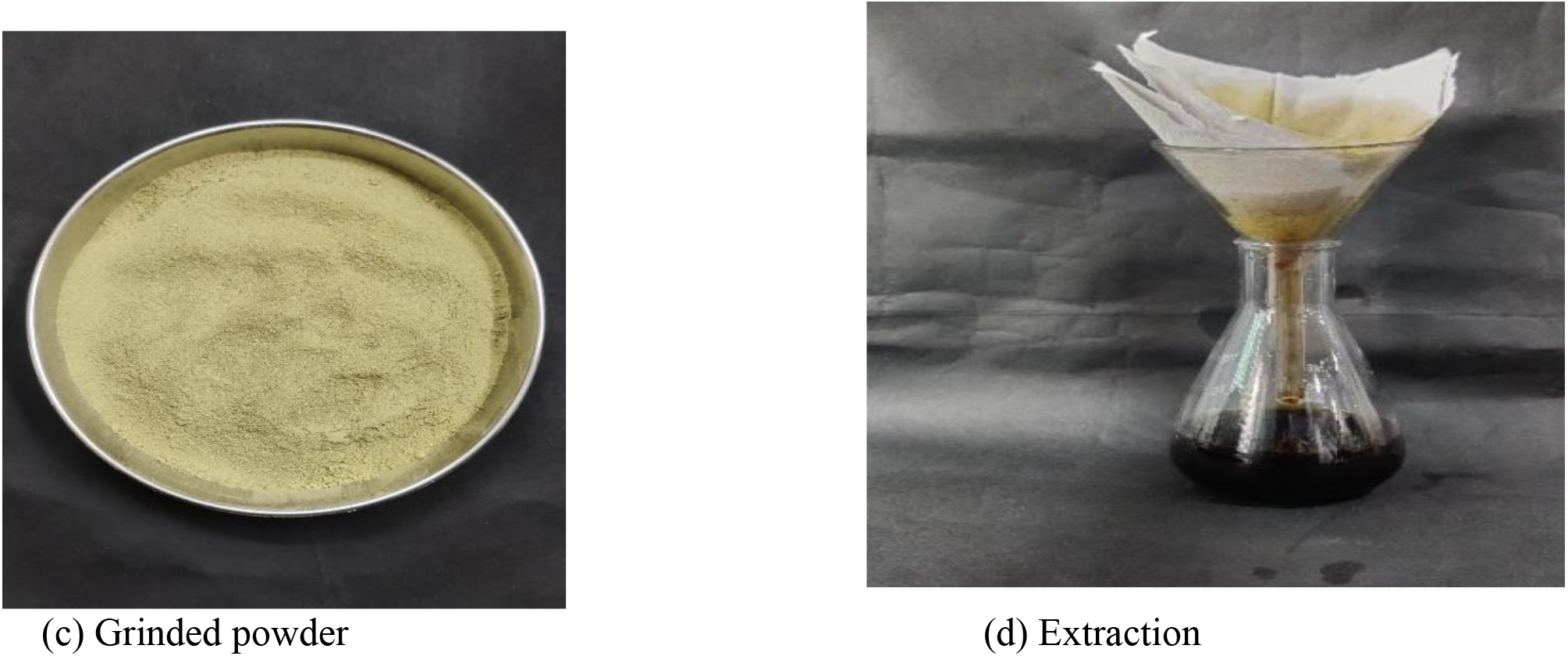
Aloe vera

### 2.2 Dinitro Salicylic Acid Test

DNSA solution has been prepared with 10 g of potassium tartrate was dissolved in 30 mL of distilled water (Solution A), and 1.5 g of dinitro salicylic acid (DNSA) was dissolved in 30 mL of sodium hydroxide (Solution B). Solutions A and B were mixed thoroughly. Six test tubes were allocated for each extract type (methanol and ethanol), with varying volumes of the extracts (0.0, 0.2, 0.4, 0.6, 0.8, and 1 mL) were added to test tubes, adjusting the total volume to 1 mL with distilled water. Subsequently, 1 mL of DNS reagent was added to each test tube. The mixtures were incubated in a boiling water bath for 10 minutes, after which the optical density was taken at 670 nm [7].

### 2.3 Biuret Test

For the Biuret test, a solution was prepared in 100 ml distilled water by dissolving 1g of copper (II) sulphate (CuSO_4_) and 1.2 g of sodium potassium tartrate (Solution 1). A second solution was made in 90 ml distilled water by dissolving 10g of sodium hydroxide (Solution 2). 10 millilitres of sodium hydroxide solution were added to solution 1. 6 test tubes were prepared with varying volumes of the sample (0.0, 0.2, 0.4, 0.6, 0.8, and 1 mL), with the total volume adjusted to 1 mL using distilled water. 3 millilitres of Biuret reagent were added in to test tubes, and mixtures were incubated at room temperature for five minutes and the optical density was taken at 580 nm [8].

### 2.4 Barfoed’s Test

For the Barfoed’s test standard preparation, a stock solution was made by dissolving 0.1g of glucose in 10 mL of distilled water. In five separate test tubes, varying volumes (0.2, 0.4, 0.6, 0.8, and 1mL) of glucose solution were added, with distilled water used to make up the total volume to 1mL in each tube. 2 millilitres of Barfoed’s reagent were then added to each test tube, which were subsequently placed in a boiling water bath for five minutes. For the sample analysis, five new test tubes were prepared with varying volumes (0.2, 0.4, 0.6, 0.8, and 1 mL) of the sample solution, again making up the volume to a total of 1 mL with distilled water in each tube and adding Barfoed’s reagent as before. After boiling for five minutes, the optical density was measured at 540nm and compared against the standard glucose solutions [9].

### 2.5 Qualitative Test for Steroids

The qualitative test for steroids was conducted, 0.2 mL of the sample was mixed with 2mL of concentrated sulphuric acid (H_2_SO_4_) plus 2 mL of chloroform. The coloured ring formation indicated the presence of steroid content [7].

## 3. Antimicrobial activity of sample

### 3.1 Preparation of sterile plates & bacterial inoculation

To make nutrient agar (100ml distilled water), take 1 conical flask, add 0.3 gm yeast extract, 0.5 gm peptone, 0.5 gm NaCl and 2 gm agar. Keep it in autoclave. Take 3 sterile petri plates, keep 1 plate as control and 2 plates for sample, pour the nutrient agar in each plate and allow it to solidify. Take *E. coli* bacteria, insert sterile inoculating loop in it and spread that bacterial culture in petri plate respectively. Repeat the same process for each plate [10].

### 3.2 Well diffusion method

The bacterial test organism E. coli was spread over the LB Agra plates by using separate sterile spreader. After 30 min at 4°C, plates were punched using cork borer. About 0.02 ml, 0.03 ml, 0.04 ml extract was delivered into each of the wells. The plates were incubated at 37°C for 24 hours, and closely monitored for the development of clear zones of inhibition around the extracts. The antimicrobial activity was assessed by the diameter of the inhibition zone.

## 4. Results

The phytochemical studies on freshly grown Aloe vera (Aloe barbadensis) are presented herein. The Aloe vera was chopped into small pieces and air-dried. The dried material was then ground to a fine powder. Ethanol and methanol extracts were prepared for subsequent testing.

### 4.1 DNSA Test

The experiment involved the preparation of methanol and ethanol extracts, which were analysed for their concentrations. The methanol extract was found to have a total concentration of 450 mg/ml, while the ethanol extract demonstrated a higher total concentration of 870 mg/ml. These findings indicate a difference in extraction efficiency of the phyto-compounds in the two solvents, with ethanol showing greater efficacy in extracting the desired substances under the given experimental conditions. The total concentration of methanol extract was 450 mg/ml and total concentration of ethanol extract was 870 mg/ml.

The test results are given in the Table-1 and Fig-2.

**Table 1.**

| Test Tubes | Sample |  | Volume of D/W (ml) | Volume of DNSA reagent | Reaction Concentration (g/ml) | OD at 670nm |  |
| --- | --- | --- | --- | --- | --- | --- | --- |
|  | Ethanol (ml) | Methanol (ml) |  |  |  | Ethanol (ml) | Methanol (ml) |
| 1. | 0.0 | 0.0 | 1 | 1 ml | 0.0 | 0.112 | 0.072 |
| 2. | 0.2 | 0.2 | 0.8 | 1 ml | 0.6 | 2.057 | 1.609 |
| 3. | 0.4 | 0.4 | 0.6 | 1 ml | 1.2 | 3.035 | 2.9 |
| 4. | 0.6 | 0.6 | 0.4 | 1 ml | 1.8 | 4.059 | 4.400 |
| 5. | 0.8 | 0.8 | 0.2 | 1 ml | 2.4 | 6.255 | 5.600 |
| 6. | 1.0 | 1.0 | 0.0 | 1 ml | 3.0 | **7.525** | **6.001** |

**Figure 2.**
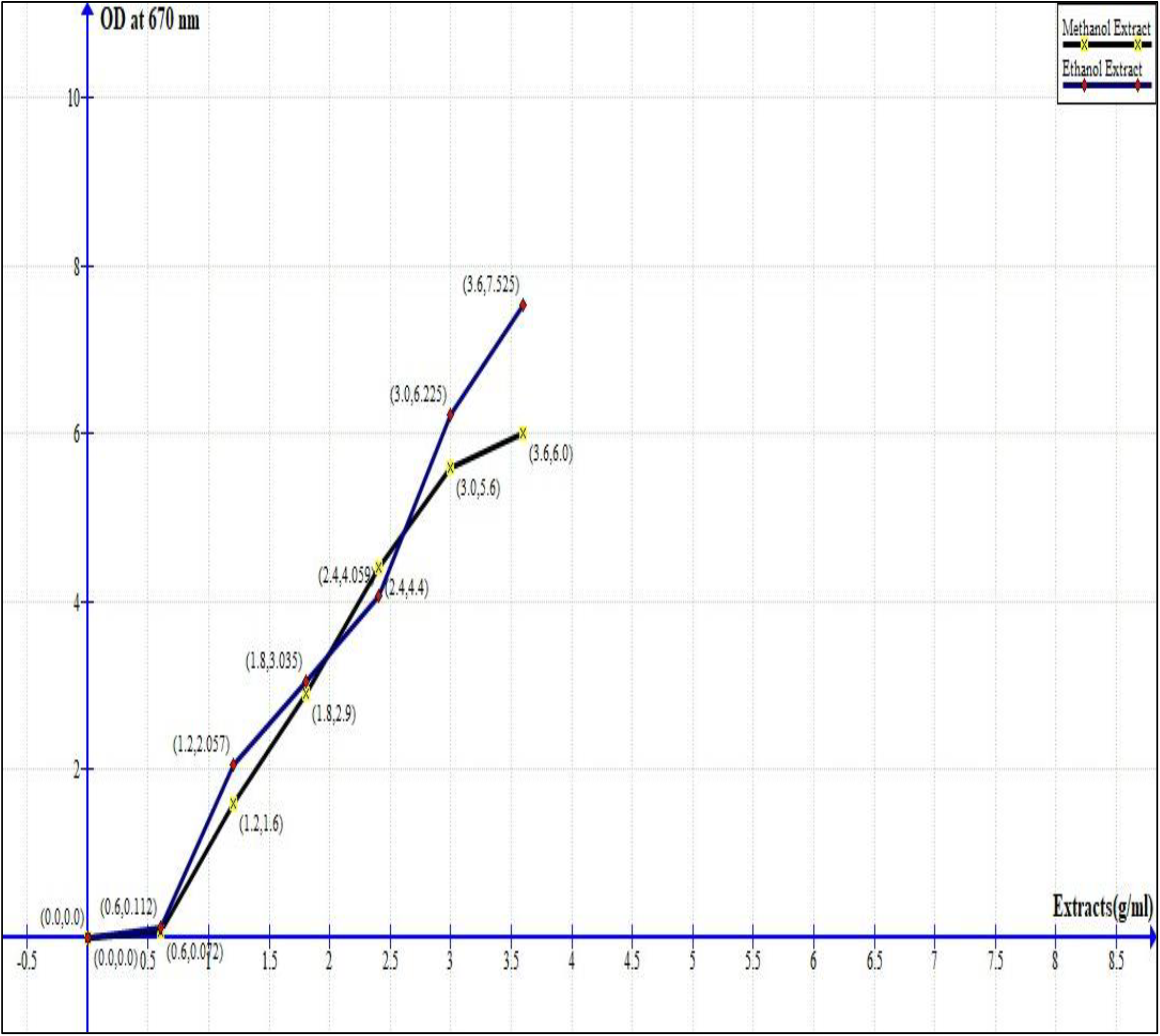

### 4.2 Biuret Test

The Biuret test was conducted to analyse the protein content in methanol and ethanol extracts. The results showed that the total concentration of the methanol extract was 13.03 mg/ml, while the ethanol extract exhibited a slightly higher total concentration of 13.8 mg/ml. The values suggest that both methanol and ethanol were effective solvents for protein extraction, with ethanol demonstrating a marginally higher efficiency under the given experimental conditions. Total concentration of methanol extract was 13.03 mg/ml and total concentration of ethanol extract was 13.8 mg/ml respectively. The test results are given in the Table-2 and Figure-3.

**Table 2.**

| Test Tubes | Sample |  | Volume of D/W (ml) | Volume of Biuret reagent | Reaction Concentration (g/ml) | OD at 540 nm |  |
| --- | --- | --- | --- | --- | --- | --- | --- |
|  | Ethanol (ml) | Methanol (ml) |  |  |  | Ethanol (ml) | Methanol (ml) |
| 1. | 0.0 | 0.0 | 1 | 3 ml | 0.0 | 1.852 | 1.922 |
| 2. | 0.2 | 0.2 | 0.8 | 3 ml | 0.6 | 3.295 | 3.298 |
| 3. | 0.4 | 0.4 | 0.6 | 3 ml | 1.2 | 3.677 | 3.510 |
| 4. | 0.6 | 0.6 | 0.4 | 3 ml | 1.8 | 3.803 | 3.850 |
| 5. | 0.8 | 0.8 | 0.2 | 3 ml | 2.4 | 4.186 | 4.138 |
| 6. | 1.0 | 1.0 | 0.0 | 3 ml | 3.0 | **5.374** | **4.359** |

**Figure 3.**
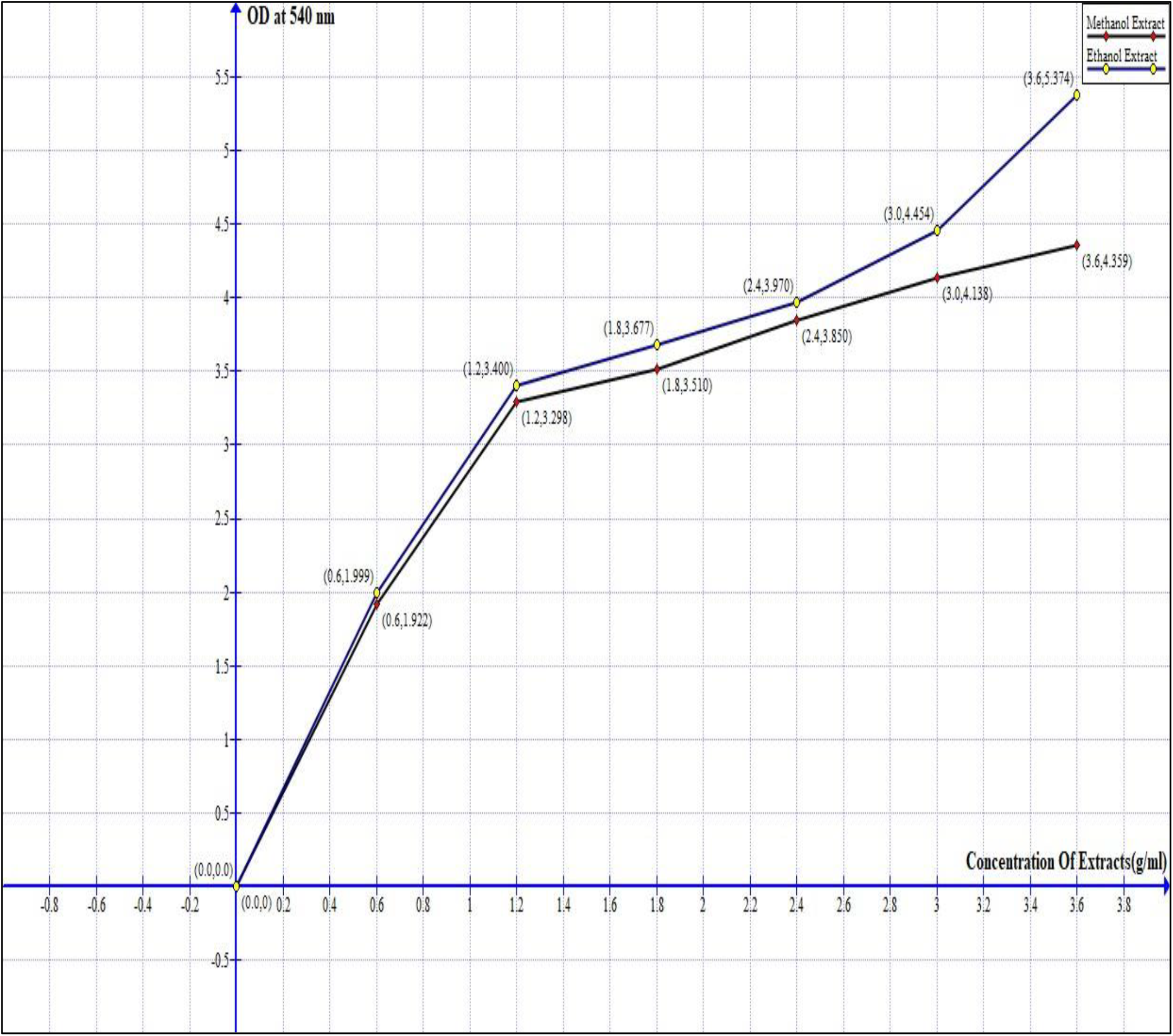

### 4.3 Barfoed’s Test

The Barfoed’s test was performed to detect the presence of monosaccharides in the sample. The analysis revealed a total concentration of 960 mg/ml, indicating a significant presence of monosaccharides in the extract under the given experimental conditions. The result highlights the efficiency of the extraction method in isolating monosaccharide components. The test results are given in the Table-3 and Figure-4.

**Table 3.**

| Test Tubes | Sample |  | Volume of D/W (ml) | Volume of Barfoed reagent | Reaction Concentration (mg/ml) | OD at 540 nm |  |
| --- | --- | --- | --- | --- | --- | --- | --- |
|  | Standard Glucose (ml) | Ethanol (ml) |  |  |  | Standard Glucose (ml) | Ethanol (ml) |
| 1. | 0.2 | 0.2 | 0.8 | 2 ml | 20 | 0.478 | 2.017 |
| 2. | 0.4 | 0.4 | 0.6 | 2 ml | 40 | 0.510 | 2.940 |
| 3. | 0.6 | 0.6 | 0.4 | 2 ml | 60 | 0.515 | 3.700 |
| 4. | 0.8 | 0.8 | 0.2 | 2 ml | 80 | 0.524 | 4.175 |
| 5. | 1 | 1 | 0.0 | 2 ml | 100 | **0.689** | **4.533** |

**Figure 4.**
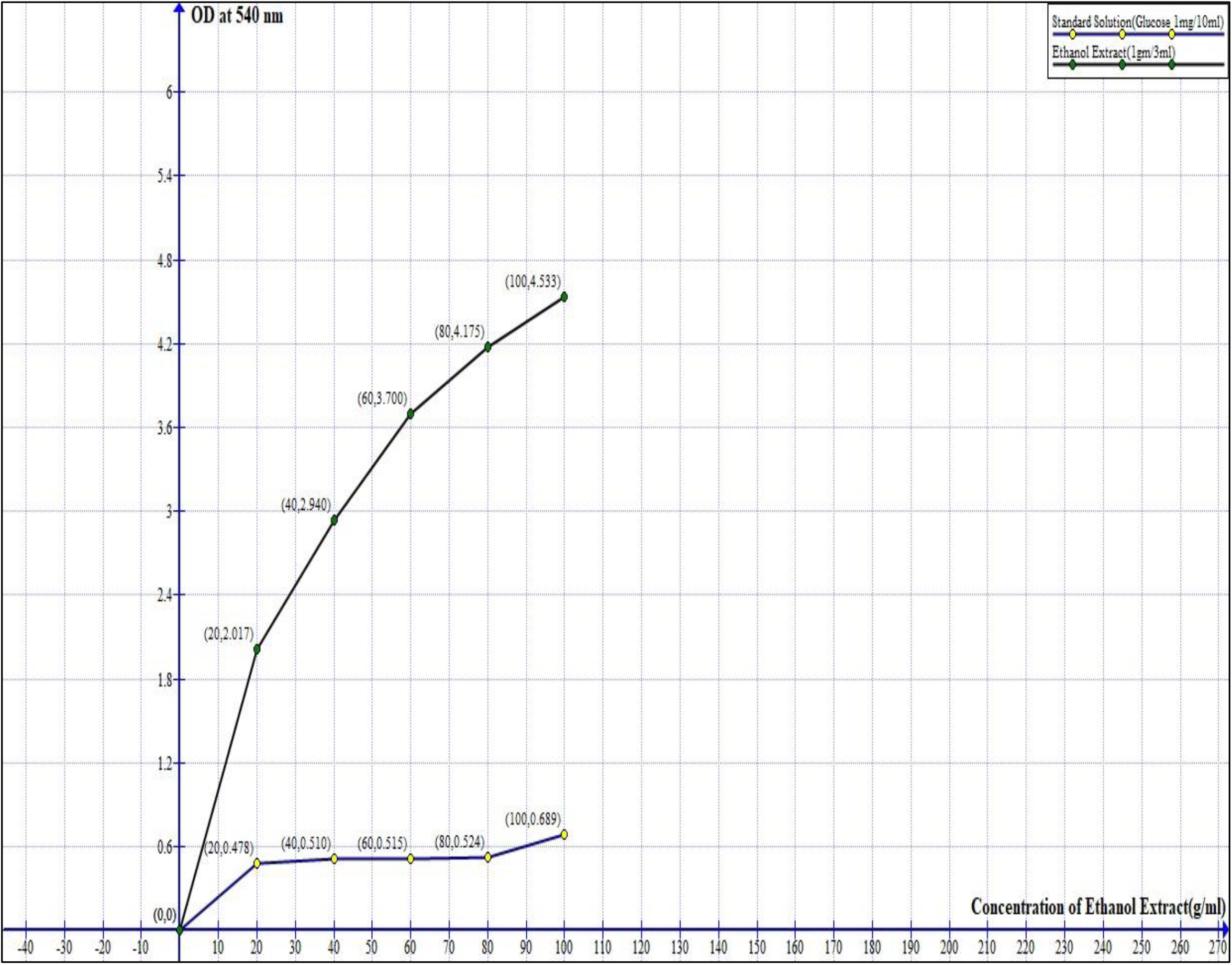

### 4.4 Qualitative Test for Steroids

The qualitative test for steroids was conducted by mixing 0.2 mL of the sample with 2 mL sulphuric acid (H_2_SO_4_) and 2 mL of chloroform. Upon addition of the reagents, a colour change was observed, indicating the steroid compounds in the sample. Specifically, the formation of a coloured ring at the interface of the sulphuric acid and chloroform layers confirmed the presence of steroid content. The result suggests that the tested Aloe vera extract contains steroidal compounds, which may contribute to its potential therapeutic properties.

### 4.5 Antimicrobial Activity

The results of the zone of inhibition measurements show varying antimicrobial effects of E. coli based on the volume of the sample used. In a 20 µL volume, the zone of inhibition measured 8 mm indicating a moderate antimicrobial effect. When the volume was increased to 40 µL, the zone of inhibition expanded to 12 mm, demonstrating a stronger antimicrobial activity with the larger volume. However, when the volume was set to 30 µL, the zone of inhibition decreased slightly to 7mm. The decrease size suggests the relationship between sample volume and antimicrobial effect is not entirely linear. The variation could be due to factors such as the concentration of the antimicrobial agent reaching a saturation point or differences in how the sample diffuses through the medium. Overall, the largest zone of inhibition was observed with the 40 µL volume indicates a higher volume tends to produce a greater antimicrobial effect of E coli. The results of antimicrobial activity of E. coli are given (control in Fig-5e and Fig-5F).

**Fig 5e.**
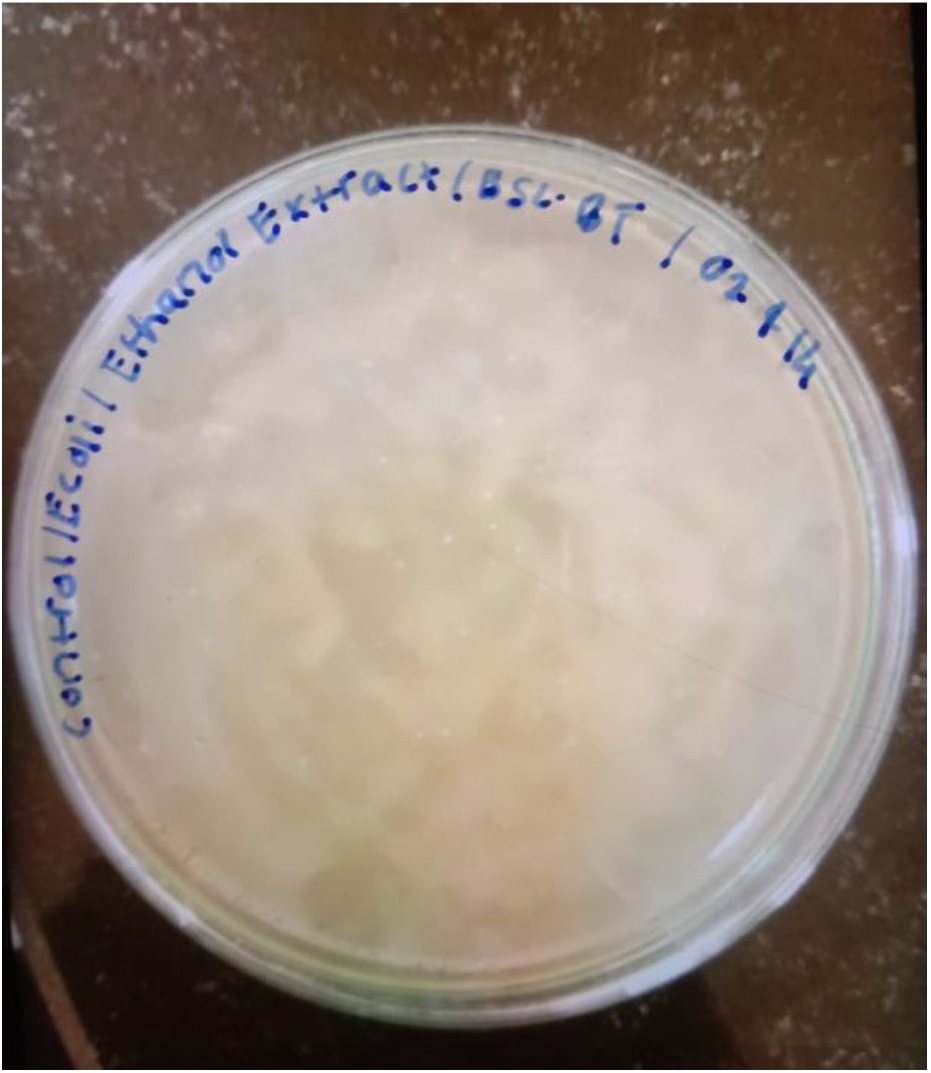
(Control, No Zone inhibition)

**Fig 5f.**
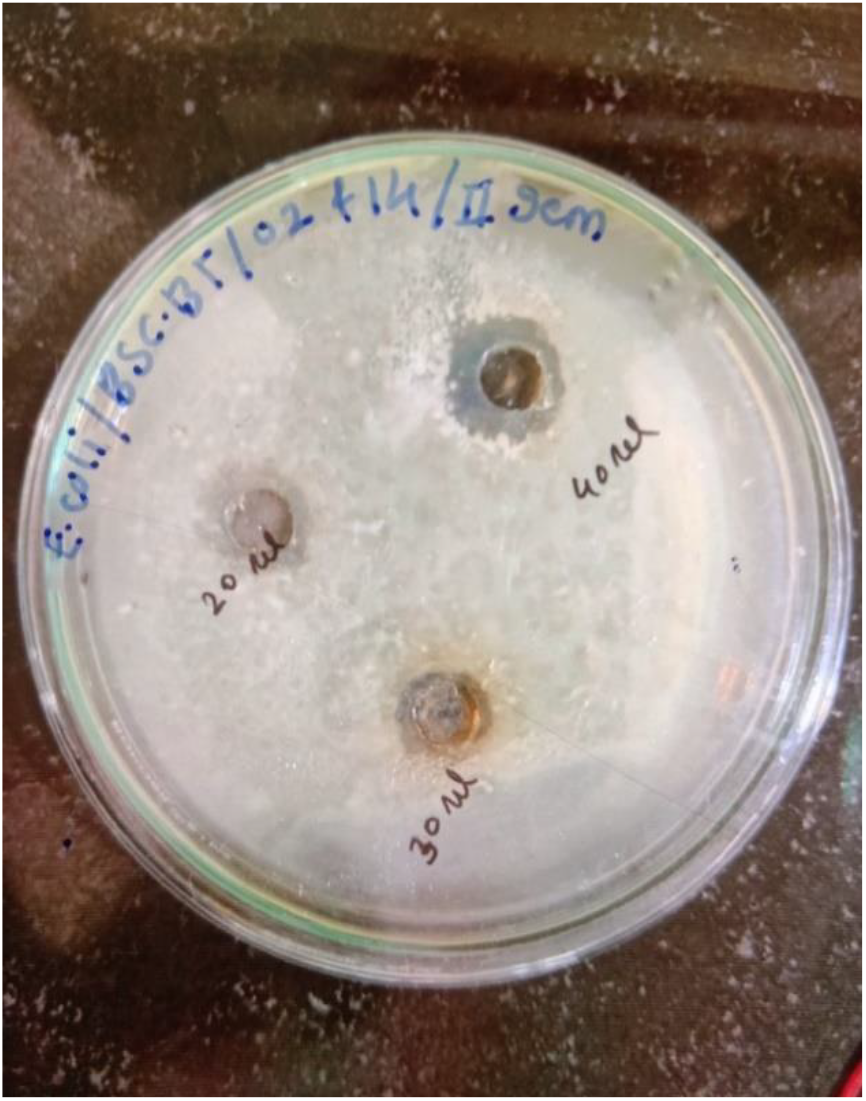
(Zone of inhibition by E coli)

## 5. Discussion

In this study, using ethanol and methanol as extraction solvents, the study examined the phytochemical characteristics of recently harvested Aloe vera (Aloe barbadensis). Plant extracts were prepared for the study and then a number of qualitative and quantitative tests were conducted to determine whether different phytochemicals, such as proteins, carbohydrates, and steroids are present. Although the results of the qualitative test for steroids showed that both extracts contained steroid components, the ethanol extract seemed to have a greater concentration of these chemicals than the methanol extract. Furthermore, the DNSA and Biuret test findings showed that ethanol was superior than methanol in terms of its ability to extract proteins and carbohydrates from Aloe vera. The results demonstrate well for ethanol to works as a solvent to extract important phytochemicals from Aloe vera.

## 6. Conclusion

Phytochemical content analysis is an important study for the Aloe barbadensis obtained from Marathwada region, India. Various phytochemical contents were analysed through a suitable ethanol extracted method and performed an antimicrobial test.

## Acknowledgements

Thanks to Administration, MGM University, Chhatrapati Sambhajinagar, Maharashtra, India. Thanks to Governor Gretchen Whitmer, Michigan State, USA and Governor Kathy Hochul, New York, USA. Thanks to The Obama Foundation, USA (www.Obama.org).

